# Visual attention is not always spatially coupled to subsequent oculomotor program

**DOI:** 10.1101/216572

**Authors:** Luca Wollenberg, Heiner Deubel, Martin Szinte

## Abstract

The premotor theory of attention postulates that spatial attention arises from the activation of saccade areas and that the deployment of attention is the consequence of motor programming. Yet, attentional and oculomotor processes have been shown to be dissociable at the neuronal level in covert attention tasks. To investigate a potential dissociation at the behavioral level, we instructed human participants to saccade towards one of two nearby, competing saccade cues. The spatial distribution of visual attention was determined using oriented Gabor stimuli presented either at the cue locations, between them or at several other equidistant locations. Results demonstrate that accurate saccades towards one of the cues were associated with presaccadic enhancement of visual sensitivity at the respective saccade endpoint compared to the non-saccaded cue location. In contrast, averaging saccades, landing between the two cues, were not associated with attentional facilitation at the saccade endpoint, ruling out an obligatory coupling of attentional deployment to the oculomotor program. Rather, attention before averaging saccades was equally distributed to the two cued locations. Taken together, our results reveal a spatial dissociation of visual attention and saccade programming. They suggest that the oculomotor program depends on the state of attentional selection before saccade onset, and that saccade averaging arises from unresolved attentional selection.

## Introduction

To process information from our rich visual environment, we evolved with attentional mechanisms allowing us to discriminate which flow to account for from which to ignore (Carrasco, 2011; Treue, 2001). For example we can extract salient saccade targets from a cluttered visual scene to later examine their contents with precise foveal vision (Adeli, Vitu, & Zelinsky, 2017; Baluch & Itti, 2011; Fecteau & Munoz, 2006; Itti & Koch, 2001). This link between attention and saccade led researchers to propose that spatial visual attention is directly dependent on the oculomotor system (Rizzolatti, Riggio, & Sheliga, 1994; Rizzolatti, Riggio, Dascola, & Umiltá, 1987), introducing what they called the "premotor theory of attention".

This influential theory of visual attention relies on two main hypotheses. The first hypothesis states that visual spatial attention is operated by the oculomotor system itself. Indeed, overlapping neuronal activations have been observed in visual attention tasks involving the deployment of attention with (overt) or without (covert) eye movements in fMRI (Corbetta, 1998). These activations include cortical and sub-cortical areas such as the Frontal Eye Fields (FEF), the parietal cortex and the Superior Colliculi (SC). Furthermore, at the behavioral level, it was for example shown that attention, measured as a local improvement in visual sensitivity, is allocated to the saccade target before the eyes start to move (Deubel & Schneider, 1996; Kowler, Anderson, Dosher, & Blaser, 1995).

The second hypothesis of the premotor theory of attention implies that the deployment of visual attention is always preceded by an activation of the oculomotor system. Under this hypothesis, covert attention involves the preparation of a saccade that is canceled before the eyes move. In line with this hypothesis, sub-threshold micro-stimulation of the FEF or the SC, which did not systematically lead to a saccade, resulted in attentional benefits measured both behaviorally and electrophysiologically at the stimulated movement field position (Juan, Shorter-Jacobi, & Schall, 2004; McPeek & Keller, 2004; Moore & Armstrong, 2003; Moore & Fallah, 2004; Müller, Philiastides, & Newsome, 2005). However, as micro-stimulation effects cannot be solely restricted to the motor cells within the stimulated areas, these results did not demonstrate that the deployment of visual attention is preceded by a premotor activation alone. Instead, it was shown that motor cells within FEF or SC stayed completely silent during a covert attention task (Gregoriou, Gotts, & Desimone, 2012; Ignashchenkova, Dicke, Haarmeier, & Thier, 2004; Thompson, Biscoe, & Sato, 2005), while visual and visuomotor cells displayed sustained attentional effects. In other words, attention is not always preceded by a motor activity, at least not within these recorded oculomotor centers. To test this second hypothesis at the behavioral level, one can imagine measuring visual sensitivity at the intended saccade goal and at the endpoint of the saccade. Under such conditions, measured sensitivity should correlate with the activity of the visual and motor cells within oculomotor centers, respectively. Taking advantage of the fact that saccades tend to undershoot the target, Deubel and Schneider (1996) found that attention was restricted to the intended saccade goal rather than to the saccade endpoint. However, using saccadic adaptation to decrease the saccadic gain, some authors found the exact opposite effect, with attention allocated to the adapted saccade endpoint rather than to the intended saccade goal (Collins & Doré-Mazars, 2006; Doré-Mazars & Collins, 2005).

Knowing that oculomotor centers have several overlapping large receptive fields within the range of these effects (Bruce & Goldberg, 1985; Schiller & Koerner, 1971), it is hard to link these contradictory behavioral findings to the neurophysiology described above. It could be interesting to use a paradigm leading to a larger spatial dissociation between the intended saccade goal and the saccade endpoint, such as the global effect (Coren & Hoenig, 1972; Findlay, 1982; Van der Stigchel & Nijboer, 2011; Vitu, 2008). Indeed, the global effect is associated with systematic and large saccade endpoint deviations towards the center of gravity of two saccade targets (Deubel, Wolf, & Hauske, 1984; Findlay, 1982; Ottes, Van Gisbergen, & Eggermont, 1984), or of a saccade target and a distractor (Ottes, Van Gisbergen, & Eggermont, 1985; Walker, Deubel, Schneider, & Findlay, 1997), shown at two positions separated by up to 60° of rotation (Ottes et al., 1985). Although the global effect was originally described as reflecting a low-level averaging of neuronal activity (and therefore respective saccades are often called averaging saccades) within the oculomotor centers (Becker & Jürgens, 1979; Findlay, 1982; Ottes, Van Gisbergen, & Eggermont, 1986), different behavioral observations later suggested a dependency on higher-level attentional processes. First, it was shown that averaging saccades can be elicited by second and third order saccade targets (Deubel, Findlay, Jacobs, & Brogan, 1988; Findlay, Brogan, & Wenban-Smith, 1993), suggesting that the global effect cannot merely reflect low-level oculomotor processes. Next, it was shown that specifying the location (Aitsebaomo & Bedell, 2000; Coëffé & O’Regan, 1987), the identity (Heeman, Theeuwes, & Van der Stigchel, 2014; Silvis, Solis, & Donk, 2015) or the probability of a saccade target to appear at a certain location relative to a distractor (He & Kowler, 1989), systematically reduced the occurrence of averaging saccades. Interestingly, monkeys make averaging saccades when the FEF or the SC are simultaneously micro-stimulated at two sites (Robinson, 1972; Robinson & Fuchs, 1969; Schiller, True, & Conway, 1979) or when two targets are shown in close proximity (Arai, McPeek, & Keller, 2004; Chou, Sommer, & Schiller, 1999). At the neuronal level, it was first proposed that a single peak of motor cell activity associated with saccades ending in between two targets precedes an averaging saccade (Glimcher & Sparks, 1993; Van Opstal & Van Gisbergen, 1990). Later work suggested instead that averaging saccades follow two peaks of activity associated with saccades directed towards the two saccade targets (Edelman & Keller, 1998; Port & Wurtz, 2003). Recently, Vokoun and colleagues (Vokoun, Huang, Jackson, & Basso, 2014) used voltage imaging of slices of rat SC to record population dynamics in response to dual-site electrical stimulation. They observed that the simultaneous stimulation of two nearby sites in the intermediate layers lead to a merged peak centered in between them in the superficial layers. Moreover, they propose that such merged activation feeds back into the visual system, leading to the perception of a target at the averaging saccade endpoint.

If this proposal of a feedback of merged activation from the superficial layers of the SC into the visual system was true, we would expect to find a presaccadic enhancement of attention at the endpoint of averaging saccades, a result that would be in line with the premotor theory of attention. Van der Stigchel and de Vries (2015) directly tested this proposal, instructing participants to move their eyes towards a saccade target presented simultaneously with a distractor and measuring presaccadic attention at these positions as well as in between them. They observed both averaging saccades as well as saccades directed towards the target and the distractor, allowing them to compare the deployment of attention at the intended saccade goal and at the saccade endpoint. Unfortunately, they report no main effect of the saccade landing direction as well as no interaction between the saccade landing direction and the position of their attention probes when they analyzed visual discrimination performance as a function of the saccade endpoint. Therefore, contrary to many reports of presaccadic attention (Deubel & Schneider, 1996; Kowler et al., 1995), the saccade landing position did not significantly affect the deployment of attention in their target-distractor paradigm, preventing any conclusion at that stage.

Various other studies have argued that visual attention spreads over an extended region in space when saccades are executed towards the center of gravity within a stimulus configuration (Cohen, Schnitzer, Gersch, Singh, & Kowler, 2007; e.g. He & Kowler, 1989; Kowler et al., 1995; Vishwanath & Kowler, 2003). Yet, none of these studies measured visual attention at the endpoint of averaging saccades. Here, we measured visual attention at various locations in space, including the averaging saccade endpoint, in a free choice saccade task which entailed the presentation of two nearby saccade targets. Our design therefore allowed us to investigate whether attention is mandatorily coupled to the saccade endpoint and therefore deployed at the endpoint of averaging saccades. Moreover, given the spatial resolution of our design, we could disentangle whether attention indeed spreads over an extended region in space or whether it is selectively deployed at the competing saccade targets. We observed a presaccadic enhancement of visual sensitivity at the endpoint of accurate saccades (landing at one of the saccade targets) but not averaging saccades (landing in between the two saccade targets), ruling out an obligatory coupling of attention to the subsequent oculomotor program. Moreover, and contrary to the idea of an extended spread of attention around the center of gravity, averaging saccades were associated with a concise and moderate enhancement of visual sensitivity at the two saccade targets. Our results therefore suggest that the oculomotor program depends on the state of attentional selection before saccade onset and that saccade averaging results from uncompleted attentional selection.

## Results

Our goal was to determine whether the presaccadic deployment of attention is coupled to the intended saccade goal or to the final saccade endpoint. To do so, we probed visual attention by presenting a discrimination target while participants prepared a saccade towards one of two potential saccade targets, presented either transiently or continuously, and separated by an inter-target angular distance of either 90° or 30° (Figure 1A). Just before the saccade, a discrimination target was shown randomly across trials at one of the two potential saccade targets (ST_1_ and ST_2_), at the position in between the saccade targets (BTW) or at one of 21 equidistant control positions (CTRL).

**Figure 1.**
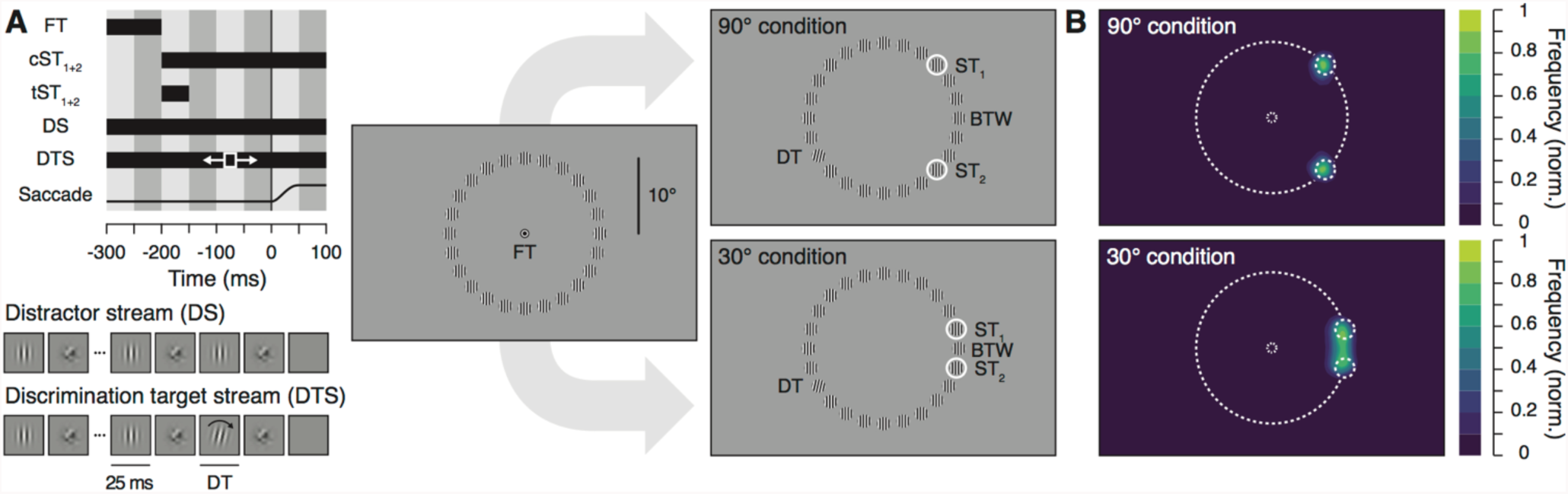
Experimental procedure and normalized saccade landing frequency maps. **A**. Stimulus timing and display. Participants prepared a saccade from the fixation target (FT) to the one of two potential saccade targets (ST_1_ and ST_2_), presented simultaneously at two randomly chosen stimuli streams with an inter-target angular distance of either 90° (top panels) or 30° (bottom panels). The saccade targets were either shown continuously (cST_1+2_) or transiently (tST_1+2_). Stimuli streams could either be distractor streams (DS), composed of alternating vertical Gabors and masks (40 Hz) or discrimination target streams (DTS) which included the presentation of a brief discrimination target (DT, 25ms), a clockwise or counter-clockwise tilted Gabor, shown between 75 and 175 ms after the saccade targets onset. Participants saccaded towards one of the saccade targets and had to report the orientation of the discrimination target, appearing randomly at one of the 24 stimuli stream locations. Note that stimuli are sketched in order to increase their visibility. Actual stimuli match those shown in the stimulus streams depiction. **B**. Normalized saccade landing frequency maps averaged across participants (n=10) for the 90° (top) and 30° (bottom) conditions (collapsed across the transient and continuous ST presentation).

Figure 1B shows the normalized frequency of saccade landing endpoints observed across participants within the 90° and 30° condition, irrespective of the duration of the saccade targets (i.e. transient and continuous combined). While saccades were equally distributed over the two saccade targets in the 90° condition (Figure 1B, top), a substantial proportion of saccades ended in between them in the 30° condition (Figure 1B, bottom). To further analyze our data, we looked at the distribution of saccade landing directions either binned in evenly distributed angular sectors of 5° (Figure 2A–B) or 15° (centered on the 24 stimuli streams, Figure 2C–D). In the 90° condition (Figure 2C), 41.0 ± 1.0 % of the saccades ended within the sector including ST_1_ (most counter-clockwise saccade target) and 41.8 ± 1.9 % within the sector including ST_2_ (most clockwise saccade target). Note that an average of 4.0 ± 0.9 % of saccades ended within the sectors adjacent to the saccade targets. In the 30° condition (Figure 2D), approximately a third (33.6 ± 2.4 %) of the saccades ended within the sector in between the two saccade targets (BTW), while 29.9 ± 1.6 % of the saccades ended within the sector of ST_1_ and 32.0 ± 1.8 % within the sector of ST_2_. Therefore, when participants had to select between two equidistant saccade targets separated by an angular distance of 30°, they executed an averaging saccade (ending in the BTW sector) in about a third of the trials. In order to determine potential differences between the two inter-target angular distance conditions (90° and 30°), we first looked at saccade latencies and amplitudes. We found slightly longer saccade latencies (90°: 192.2 ± 1.7 ms vs. 30°: 188.2 ± 2.2 ms, *p* = 0.0012) and larger amplitudes (90°: 10.0 ± 0.1° vs. 30°: 9.7 ± 0.1°, *p* = 0.0002) in the 90° as compared to the 30° condition. Saccade latency did not differ as a function of the saccade landing position (ST_1_, ST_2_ or BTW) both in the 90° and 30° condition (all *ps* > 0.05, Figure 2E–F). In the 90° condition, amplitudes of saccades towards ST_1_ (10.1 ± 0.1°) and ST_2_ (10.0 ± 0.1°), did not differ significantly from each other (ST_1_ vs. ST_2_: p = 0.7902), whereas amplitudes of saccades towards BTW (7.9 ± 0.2°) were significantly smaller than those of saccades towards ST_1_ and ST_2_ (both *ps* < 0.0001), see Figure 2G. In the 30° condition, amplitudes of saccades towards ST_1_ (9.7 ± 0.1°) and ST_2_ (9.8 ± 0.1°), as well as towards ST_1_ and BTW (9.7 ± 0.1°) did not differ significantly from each other (ST_1_ vs. ST_2_: *p* = 0.2216, ST_1_ vs. BTW: *p* = 0.5998), whereas amplitudes of saccades towards ST_2_ were significantly larger than those of saccades towards BTW (ST_2_ vs. BTW: *p* = 0.0118), see Figure 2H. Note that the proportion of averaging saccades did not vary as a function of saccade latency. Comparing trials of the 30° condition separated in two equal groups of early (167.1 ± 1.8 ms) and late (209.3 ± 3.2 ms) saccade latencies we found a comparable proportion of averaging saccades (early BTW: 35.1 ± 3.0% vs. late BTW: 32.1 ± 2.2%, *p* = 0.1632). This effect is most likely the consequence of the instruction given to the participants to saccade as fast as possible, such that early and late averaging saccade latencies differed by less than 40 ms (early BTW: 168.2 ± 2.0 ms vs. later BTW: 207.4 ± 3.1 ms, *p* < 0.0001). Overall, for each inter-target angular distance, we observed either no differences or only some non-systematic differences of a few milliseconds and a few minutes of arc. Although saccade latencies and amplitudes did not differ much between these conditions, the saccade landing direction distributions reflect two distinct oculomotor modes as a function of the inter-target angular distance. Saccades were mostly accurate in the 90° condition, whereas we observed both accurate and averaging saccades in the 30° condition.

**Figure 2.**
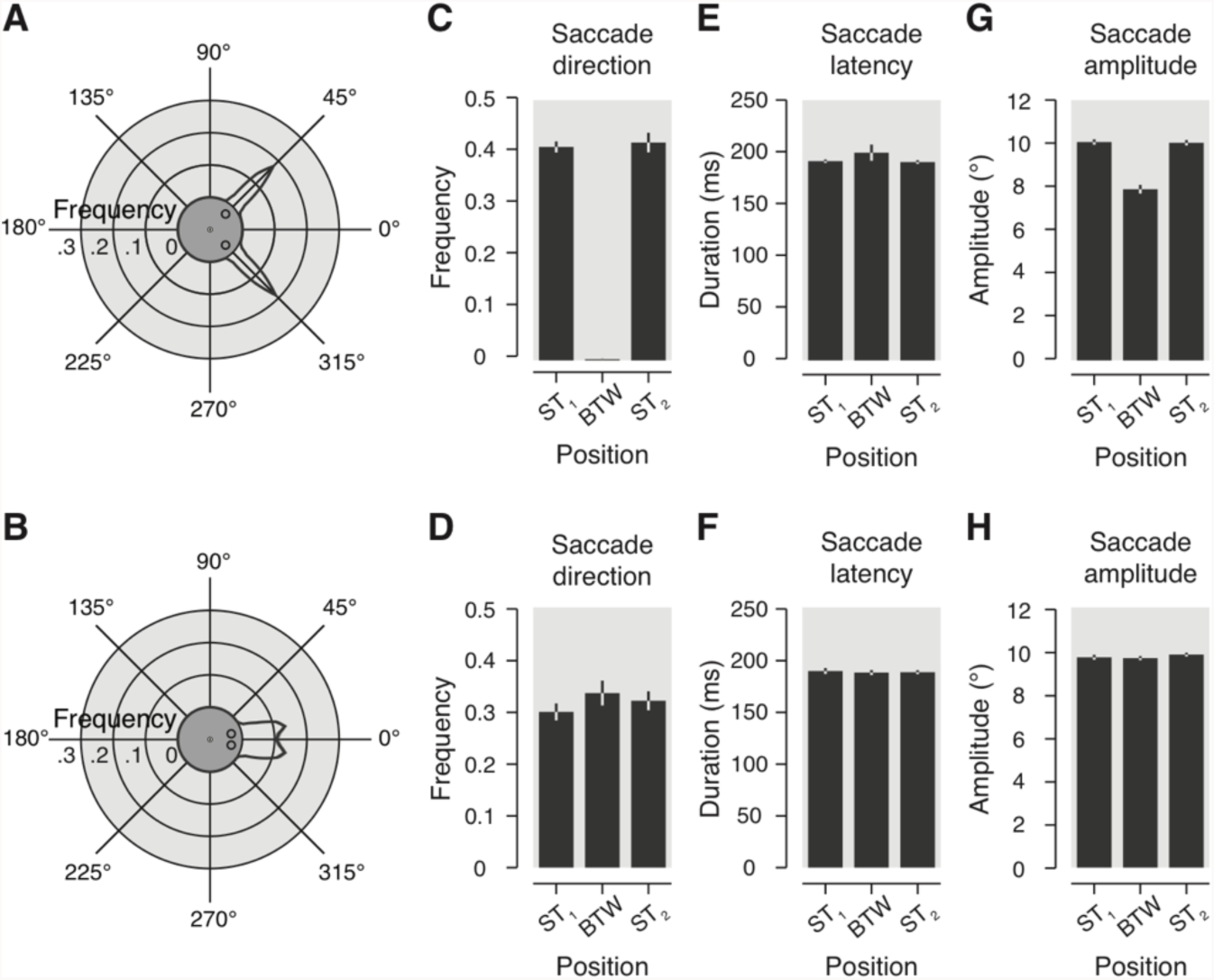
Saccade metrics. **A–B.** Circular plots show the averaged frequency distribution of the saccade landing direction binned in evenly distributed angular sectors of 5°, in the 90° (A) and 30° conditions (B). Stimuli configuration is rotated as to align the two saccade targets symmetrically around the geometrical angle zero (see central insets). **C–D.** Bar graphs illustrate averaged frequency of trials as a function of the saccade landing direction binned in 24 evenly distributed angular sectors of 15°. Data shown for the three positions of interest (ST_1_, BTW and ST_2_) in the 90° (C) and 30° conditions (D). **E–H.** Averaged saccade latency (E, F) and amplitude (G, H) observed for the same three positions of interest in the 90° (E, G) and 30° conditions (F, H). All data are shown here irrespective of the duration of the saccade targets. Light gray areas and error bars represent SEM. Polar plot black lines and corresponding light gray areas show linear interpolation between data points.

Our paradigm allowed us to measure both the oculomotor behavior and the presaccadic allocation of attention through the presentation of a discrimination target at one of 24 possible positions. We first verified that the presentation of the discrimination target itself did not systematically influence oculomotor behavior. We did not find any differences with respect to saccade latency and amplitude when comparing trials with and without the presentation of a discrimination target (3.5% of trials were without discrimination target, both *ps* > 0.05). This result validates that the distractor streams and in particular the presentation of a discrimination target did not bias the deployment of attention. Figure 3A–B show visual sensitivity as a function of the discrimination target position, rotated as to align the two saccade targets on the geometrical angle zero in both the 90° (Figure 3A) and 30° condition (Figure 3B). Irrespective of the duration of the saccade targets, we found higher sensitivity for discrimination targets shown at the saccade targets than at the control positions (corresponding to all positions except for ST_1_, ST_2_ and BTW) in both the 90° (ST_1_: d’ = 2.2 ± 0.3 vs. CTRL: d’ = 0.3 ± 0.1, *p* < 0.0001; ST_2_: d’ = 2.2 ± 0.4 vs. CTRL, *p* < 0.0001; ST_1_ vs. ST_2_: *p* = 0.8964; Figure 3A) and the 30° condition (ST_1_: d’ = 2.2 ± 0.3 vs. CTRL: d’ = 0.3 ± 0.1, *p* < 0.0001; ST_2_: d’ = 2.1 ± 0.3 vs. CTRL, *p* < 0.0001; ST_1_ vs. ST_2_: *p* = 0.6026; Figure 3B). These effects contrast with the low sensitivity observed for discrimination targets shown in between the saccade targets (BTW) in the 90° (BTW: d’ = 0.2 ± 0.1 vs. ST_1_, *p* < 0.0001; BTW vs. ST_2_, *p* < 0.0001) and especially in the 30° condition (BTW: d’ = 0.6 ± 0.2 vs. ST_1_, *p* < 0.0001; BTW vs. ST_2_, *p* < 0.0001).

**Figure 3.**
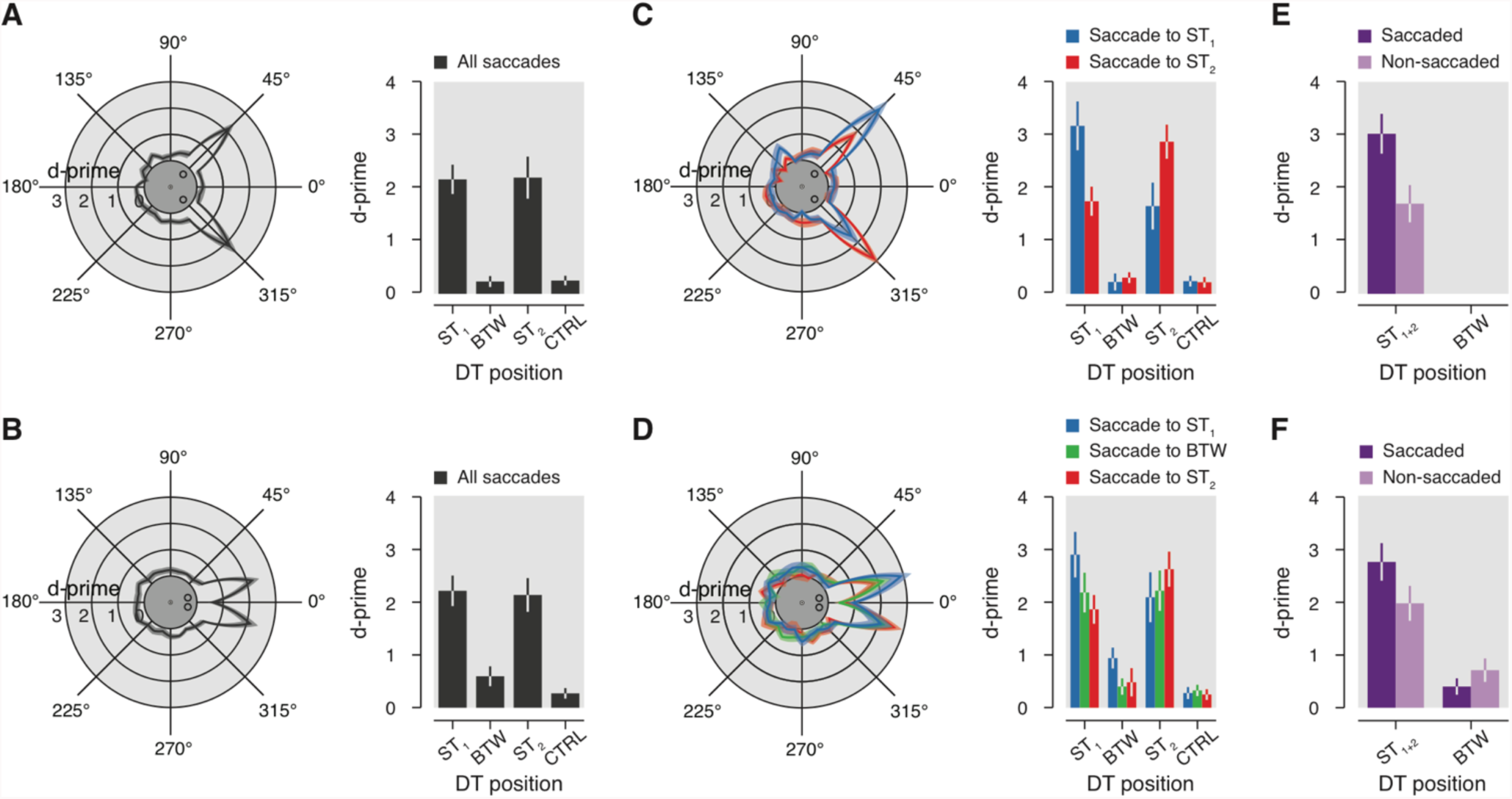
Visual sensitivity. **A–B**. Circular plots show averaged visual sensitivity (d’) as a function of the DT position in the 90° (A) and 30° conditions (B), irrespective of the duration of the saccade targets and across all saccade directions observed. Bar graphs illustrate visual sensitivity for four positions of interest (ST_1_, BTW, ST_2_, CTRL). **C–D.** Visual sensitivity as a function of the DT position relative to the saccade landing direction in the 90° (C) and 30° conditions (D), irrespective of the duration of the saccade targets (blue: saccade to ST_1_; green: saccade to BTW; red: saccade to ST_2_). For each saccade direction we took the average sensitivity for each discrimination target location. For example, the blue line plots visual sensitivity when saccades were made towards ST1 and the discrimination target was either at ST1 (+15° on the polar plot), BTW (15° counter-clockwise to ST1; 0° on the polar plot) or ST2 (30° counter-clockwise to ST1; +345° on the polar plot), and so on. **EC–F.** Bar graphs illustrate sensitivity observed for DT shown at the saccaded (purple: e.g. DT at ST_1_ and saccade to ST_1_) and the non-saccaded positions (light-purple: e.g. DT at ST_1_ and saccade to ST_2_ or BTW) in the 90° (E) and the 30° (F) conditions. Conventions are as in Figure 2.

Thus, despite the fact that saccades landed in between the saccade targets in a third of the trials in the 30° condition, the overall sensitivity at this position stayed rather low. One should however note, that sensitivity was still increased at this position compared to the control positions in the 30° condition (30°: BTW vs. CTRL, *p* = 0.0010), whereas this was not the case in the 90° condition (90°: BTW vs. CTRL, *p* = 0.7732). On the other hand, such slight facilitation observed in between the saccade targets in the 30° condition relatively to the control positions was only observed for trials in which the targets were shown transiently (BTW: d’ = 0.8 ± 0.2 vs. CTRL: d’ = 0.3 ± 0.1, *p* < 0.0001) but not continuously (BTW: d’ = 0.5 ± 0.2 vs. CTRL: d’ = 0.3 ± 0.0, *p* = 0.10880). It is important to note that the discrimination target temporally overlapped with the saccade targets in the continuous but never in the transient condition. The observed difference between the two conditions therefore suggests that the appearance of a discrimination target at BTW was masked by the continuous presentation of the saccade targets. Altogether, the results above demonstrate that presaccadic attention was mainly allocated towards the saccade targets, and to a smaller extent towards the position in between. This last result however cannot be attributed to a large spread of attention extending to more than one of the tested directions as we did not observe a consistent benefit at the two other positions adjacent to the saccade targets in the 30° condition (ST_1_ + 15°: d’ = 0.4 ± 0.1 vs. CTRL: d’ = 0.3 ± 0.1, *p* = 0.0914; ST_2_ – 15°: d’ = 0.4 ± 0.1 vs. CTRL, *p* = 0.0336; here CTRL excludes ST_1_ + 15° and ST_2_ – 15°, respectively in addition to ST_1_, ST_2_ and BTW) nor at the four adjacent positions of the saccade targets in the 90° condition (ST_1_ ± 15°: d’ = 0.3 ± 0.1 vs. CTRL: d’ = 0.2 ± 0.1, *p* = 0.5742; ST_2_ ± 15°: d’ = 0.3 ± 0.1 vs. CTRL, *p* = 0.3200; here CTRL excludes ST_1_ ± 15° and ST_2_ ± 15°, respectively in addition to ST_1_, ST_2_ and BTW).

At that stage, one cannot exclude the possibility that attention is always drawn towards the saccade endpoint in both accurate and averaging saccades, since we found higher sensitivity for both the saccade targets and in the 30° condition also for the position in between them, when compared to the control locations. Although we found higher sensitivity at the saccade targets than in between them, this may just reflect the combined effect of the saccade preparation and the presence of visual cues (the saccade targets themselves). To estimate the effect of saccade preparation, we thus needed to specify our results depending on where the saccade ended within each trial. To do so, we redefined the position of the discrimination targets relatively to the saccade direction. Figure 3C–D show visual sensitivity as a function of the discrimination target position relative to the saccade direction. We found better sensitivity for discrimination targets shown at the saccade targets when compared to the position in between them or to the control positions in both the 90° and 30° conditions, for trials in which accurate saccades were made towards ST_1_ (all *ps* < 0.0001) or ST_2_ (all *ps* < 0.0001). The same effects were found for averaging saccades in the 30° condition (all *ps* = 0.00010). In addition to the facilitation effect of the saccade targets' presentation, we found that irrespective of the inter-target distance (90° or 30°), sensitivity at ST_1_ was improved when an accurate eye movement was made towards ST_1_ (90°: ST_1_: d’ = 3.2 ± 0.5 vs. ST_2_: d’ = 1.7 ± 0.4, *p* < 0.0001, see blue lines and bars in Figure 3C–D; note that in the 30° condition, sensitivity at ST_1_: d’ = 2.9 ± 0.4 was only marginally superior to those observed at ST_2_: d’ = 2.1 ± 0.5, *p* = 0.0740). The same selective improvement was observed at ST_2_ before the execution of accurate saccades towards it (90°: ST_2_ vs. ST_1_, *p* < 0.0001; 30°: ST_2_ vs. ST_1_, *p* = 0.0002, see red lines and bars in Figure 3C–D). In particular, preparing an accurate eye movement towards one of the two saccade targets improved sensitivity when comparing trials where a discrimination target was shown at the saccaded location (e.g. DT at ST_1_ and saccade made towards ST_1_) to trials where a discrimination target was shown at the same position that was not the saccaded position (e.g DT at ST_1_ and saccade landing at ST_2_ or BTW) in both the 90° (Figure 3E; ST_1+2_ saccaded: d’ = 3.0 ± 0.4 vs. ST_1+2_ non-saccaded: d’ = 1.7 ± 0.4, *p* < 0.0001) and the 30° condition (Figure 3F; ST_1+2_ saccaded: d’ = 2.7 ± 0.4 vs. ST_1+2_ non-saccaded: d’ = 2.0 ± 0.3, *p* = 0.0080). Crucially for averaging saccade trials, for which the intended saccade goal (ST_1_ or ST_2_) and the saccade endpoint (BTW) were dissociated (see green lines and bars in Figure 3D), we found a rather low sensitivity for discrimination targets shown in between the saccade targets (BTW: d’ = 0.4 ± 0.2), highly reduced when compared to discrimination targets shown at the saccade targets (ST_1_: d’ = 2.2 ± 0.4 and ST_2_: d’ = 2.2 ± 0.4, both *ps* < 0.0001), and contrary to above (Figure 3B), not different from the sensitivity gathered across the control locations (CTRL: d’ = 0.3 ± 0.1, *p* = 0.4026), both when the saccade targets were shown transiently or continuously (both *ps* > 0.05). Thus, contrary to accurate saccades, the execution of averaging saccades did not lead to any improvement at the saccade endpoint. Rather, we found that preparing an average saccade significantly reduced participants’ sensitivity when comparing trials with discrimination targets shown at the saccaded direction to trials where the discrimination targets were shown at the same direction, which however was not the saccaded one (Figure 3F; BTW saccaded: d’ = 0.4 ± 0.2 vs. BTW non-saccaded: d’ = 0.7 ± 0.2, *p* < 0.0001). This sensitivity reduction can however mainly be attributed to a masking effect of the continuous presentation of the saccade targets (BTW saccaded: d’ = 0.3 ± 0.3 vs. BTW non-saccaded: d’ = 0.7 ± 0.2, *p* = 0.0088) as it was not found for saccade targets presented transiently (BTW saccaded: d’ = 0.7 ± 0.2 vs. BTW non-saccaded: d’ = 0.7 ± 0.3, *p* = 0.9664).

Therefore, we here demonstrated, contrary to what is predicted by the premotor theory of attention, that the preparation of averaging saccades did not lead to a deployment of attention at the corresponding saccade endpoint. Instead, we found that averaging saccades were associated with an equal distribution of attention towards the two saccade targets (ST_1_: d’ = 2.2 ± 0.4 vs. ST_2_: d’ = 2.2 ± 0.4, *p* = 0.8402). One interpretation of these effects could be that averaging saccades result from an unsuccessful or at least uncompleted presaccadic attentional selection among the two saccade targets, with resources equally distributed over them. On the other hand, it is possible that, despite landing in between the targets, presaccadic attentional selection was successful before averaging saccades but directed half of the time towards the most clockwise saccade target and half of the time towards the most counter-clockwise saccade target. If this was the case, across trials, one would also expect to find an equal and moderate enhancement of sensitivity for discrimination targets shown at the saccade targets.

To disentangle these two interpretations, we analyzed trials in which a corrective saccade followed the execution of an averaging saccade. We reasoned that if averaging saccades resulted from a successful trial-by-trial presaccadic attentional selection of one of the two saccade targets, they should be followed by corrective saccades directed equally often towards one of the targets. Moreover, they should be associated with an attentional benefit at the goal of the corrective saccade. Contrary to these predictions, we observed corrective saccades in only 48.1 ± 5.8 % of the averaging saccade trials. Corrective saccades were not all clearly directed towards the saccade targets (Figure 4A–B), ending either in the angular sector of the most counter-clockwise saccade target (ST_1_: 48.3 ± 3.1 % of all the corrective saccades following an averaging saccade), the most clockwise saccade target (ST_2_: 38.3 ± 2.5 %) or in between them (BTW: 11.9 ± 2.8 %). They were, moreover, not equally often directed towards each of the saccade targets (ST_1_ vs. ST_2_, *p* = 0.0288), probably reflecting a bias of our participants. Importantly, as shown in Figure 4C, we did not find any significant benefit at the endpoint of the corrective saccades following an averaging saccade, when comparing trials where discrimination targets were shown at the endpoint of the corrective saccade (ST_1+2_ correctively saccaded: d’ = 2.8 ± 0.5) to trials where a discrimination target was shown at the same position that was not the endpoint of the corrective saccade (ST_1+2_ correctively non-saccaded: d’ = 2.5 ± 0.8, *p* = 0.68300). Moreover, no significant benefit could be found when the corrective saccades following an averaging saccade ended still in between the saccade targets (BTW correctively saccaded: d’ = 0.7 ± 1.1 vs. BTW correctively non-saccaded: d’ = −0.1 ± 0.6, *p* = 0.4698). Taken together, these results therefore suggest that averaging saccades result from an unsuccessful or uncompleted presaccadic attentional selection among the two saccade targets.

**Figure 4.**
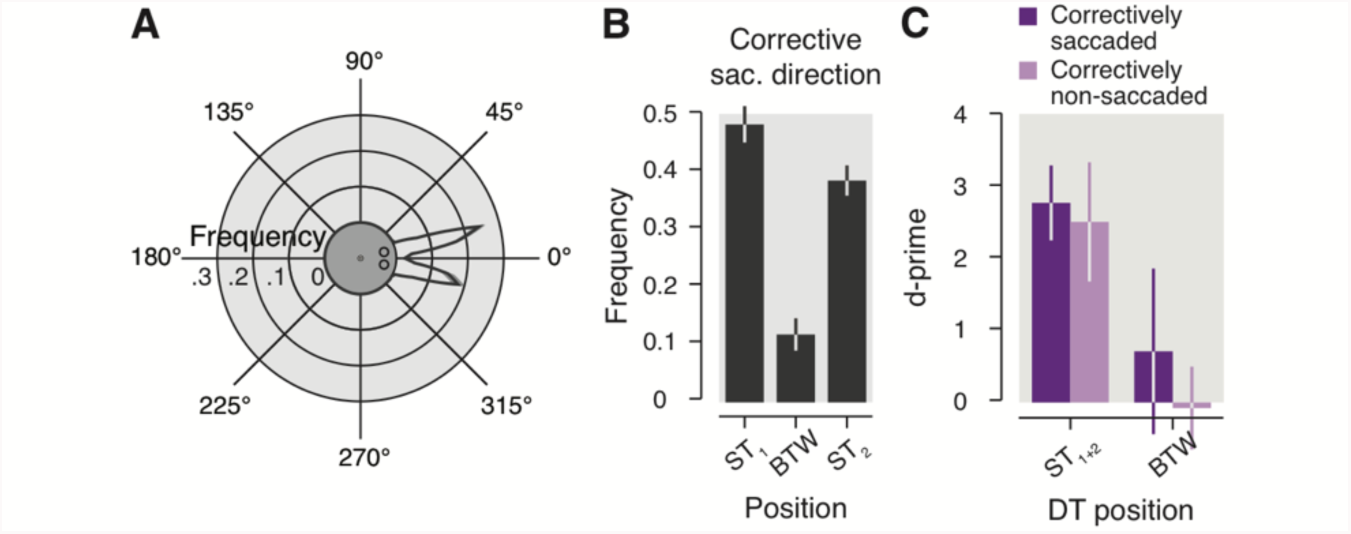
Corrective saccades. **A** Circular plot shows averaged frequency distribution of the corrective saccade landing direction following an averaging saccade. **B** Bar graph illustrates averaged frequency of trials as a function of the corrective saccade landing direction following an averaging saccade for three positions of interest (ST_1_, BTW and ST_2_). **C.** Bar graph illustrates sensitivity observed for DT shown at the correctively saccaded (purple) and the correctively non-saccaded positions (light-purple) for trials in which the main saccade was directed in between the saccade target. Conventions are as in Figure 2–3.

## Discussion

We observed a clear oculomotor dissociation between trials in which two equidistant saccade targets were shown at two different angular distances from each other. While only accurate saccades were found for an inter-target angular distance of 90°, we observed both accurate and averaging saccades when the same targets were separated by 30°. Combined with a measure of presaccadic visual sensitivity, this dissociation allowed us to determine the influence of saccade preparation on the deployment of attention when the intended saccade goal and the saccade endpoint were spatially associated (accurate saccades) or clearly dissociated from each other (averaging saccades). Accurate saccades were associated with a strong and systematic presaccadic enhancement of visual sensitivity at the saccade endpoint when compared to the non-saccaded locations for inter-target angular distances of both 90° and 30°. In contrast, averaging saccades were not associated with an attentional facilitation at the saccade endpoint. These effects rule out an obligatory coupling of visual attention to the activation of the oculomotor system and reflect a spatial dissociation between visual attention and saccade programming before averaging saccades. Interestingly, averaging saccades were associated with an equal deployment of attention at the two saccade target locations. Our corrective saccade analysis indicates that this result cannot be explained by a trial-by-trial presaccadic attentional selection of one of the two saccade targets. Therefore, contrary to the idea that the activation of the oculomotor system precedes spatial attention, we propose that the oculomotor program depends on the state of attentional selection before the saccade onset, with averaging saccades arising from an uncompleted attentional selection.

Findlay (1982) referred to the "global effect" as the phenomenon of directing the eyes towards the center of gravity of two presented targets (Coren & Hoenig, 1972). To his view, this phenomenon reflects a coarse or global processing of a visual scene before a rapidly generated eye movement. His account thus predicts that in our experiment, visual sensitivity should be coarsely distributed over the two saccade targets as well as over their adjacent locations before the execution of averaging saccades. Similarly, various other studies have argued that visual attention spreads over an extended region in space when saccades are executed towards the center of gravity within a stimulus configuration (Cohen et al., 2007; e.g. He & Kowler, 1989; Kowler et al., 1995; Vishwanath & Kowler, 2003). Our precise measure of presaccadic visual sensitivity allowed us to determine the spatial specificity of attentional deployment during saccade preparation. Contrary to the notion of a global processing mode before averaging saccades and an extended attentional spread around the center of gravity in response to global saccade target configurations (Cohen et al., 2007; e.g. He & Kowler, 1989; Kowler et al., 1995; Vishwanath & Kowler, 2003), we observed a precise allocation of attention limited only to the saccade targets (limited to a distance below at least ~2.6°, the distance between two of our adjacent stimuli). Therefore, before an averaging saccade, the visual system indeed seems to have precise access to the saccade target configuration, reflecting an enhancement of local rather than global visual information processing (Li, Barbot, & Carrasco, 2016). Such a discontinuous deployment of attention was also found in various tasks entailing the presentation of multiple targets (Baldauf & Deubel, 2008; Dubois, Hamker, & VanRullen, 2009; e.g. McMains & Somers, 2004). Importantly, our results can also rule out other models of averaging saccades based solely on low-level oculomotor processing (Arai & Keller, 2005; Becker & Jürgens, 1979; Ottes et al., 1986). We report here that when an accurate saccade is prepared towards one of two identical saccade targets, the subsequent movement correlates with an attentional benefit at the saccade endpoint, whereas averaging saccades resulted in the absence of a selective attentional benefit at one of the two targets as well as in between them (i.e. at the saccade endpoint). In this regard, our results match with previous studies showing a reduction in the occurrence of averaging saccades when the attentional selection of the saccade goal is made more easy by specifying its location or its identity (Aitsebaomo & Bedell, 2000; Coëffé & O’Regan, 1987; He & Kowler, 1989; Heeman et al., 2014; Silvis et al., 2015). Similarly, a model relying on attentional selection could also explain that averaging saccades are less often observed in delayed saccade tasks (Coëffé & O’Regan, 1987; Jacobs, 1987), as they also give more time for the attentional selection to complete (Silvis et al., 2015). Another account of the global effect is that averaging saccades reflect a time-saving strategy (Coëffé & O’Regan, 1987), in which a fast averaging saccade followed by a correction movement allows to be faster than a deliberately delayed accurate saccade. Given that participants saccaded accurately towards one of the targets with a similar latency as found for averaging saccades in two thirds of the trials in our paradigm, our results speak against such a strategy. Although we observed some corrective saccades which ended nearby the saccade targets and therefore increased the accuracy of earlier averaging saccades, they came with a cost of about 200 ms, rendering such strategy inefficient. Moreover, if participants would have strategically planned two successive saccades (an averaging saccade followed by a corrective saccade), we would expect to find attentional benefits at both saccade endpoints as reported in sequential saccade tasks (Baldauf & Deubel, 2008; Rolfs, Jonikaitis, Deubel, & Cavanagh, 2011). Contrary to this prediction, we neither found an attentional enhancement at the endpoint of averaging saccades nor at the endpoint of corrective saccades compared to the positions not reached by corrective saccades.

Therefore, we ruled out earlier accounts of the global effect and propose that averaging saccades reflect a compromise between the dynamics of attentional selection and the instructions to move the eyes as fast as possible. Our proposal is based on the results of a combined measure of attention and averaging saccades. Similar to a previous report (Van der Stigchel & de Vries, 2015) we found an overall enhancement of visual sensitivity at the two saccade targets, when the data were not split depending on the saccade direction. In order to conclude on the deployment of attention before averaging saccades one needs to specify visual sensitivity depending on the saccade direction. Crucially, and contrary to Van der Stigchel & de Vries (2015), we indeed found an influence of the saccade direction (i.e. endpoint) on the allocation of attention when taking into account saccade direction. Within a paradigm producing both accurate and averaging saccades, we observed a presaccadic shift of attention (Deubel & Schneider, 1996; Kowler et al., 1995), reflected by selectively enhanced sensitivity at the endpoint of accurate saccades. The replication of this presaccadic attention effect comes as a pre-requisite to conclude on the effect of averaging saccades, for which instead, we found no attentional benefit at the saccade endpoint. Van der Stigchel & de Vries (2015) concluded that there is no attentional shift towards the endpoint of averaging saccades. However, they also reported no main effect of the saccade landing direction as well as no interaction between the saccade landing direction and the position of their attention probes when they analyzed their data as a function of the saccade endpoint. Their results are therefore inconclusive, or even speak in favor of an attentional global effect. Moreover, when we combined all trials irrespective of the saccade direction, we found a slight increase of sensitivity at the position in between the two potential saccade targets when they were presented transiently but not when they were presented continuously. As Van der Stigchel & de Vries (2015) used a continuous presentation of a saccade target and a distractor, their results most likely reflect a masking effect of their stimuli on the discrimination target rather than an absence of attentional modulation. We here clearly dissociated attention allocated to the intended saccade goal from attention allocated to the endpoint of the saccade, and found no benefit at the averaging saccade endpoint. Consequently, our results rule out the main hypothesis of the premotor theory of attention, which postulates that the deployment of visual attention is always preceded by an activation of the oculomotor system (Rizzolatti et al., 1987; 1994).

We illustrate our results in a theoretical framework (Figure 5), inspired by both behavioral and neurophysiological findings, linking visual attention and oculomotor programming (Rolfs & Szinte, 2016). This theoretical framework does neither provide a strict model nor a computational framework. It aims at putting our results in the context of the current view on saccade programming and yielding new testable predictions. We propose that our attentional effects rely on top-down modulation (Baluch & Itti, 2011; Moore & Armstrong, 2003) of feature selective areas of the visual cortex by the priority maps (Bisley & Goldberg, 2010). Initially, the onsets of the saccade targets strongly activate neurons with corresponding receptive fields within columns of the feature and priority maps (Figure 5A). Their activity will then decay until the saccade target selection process begins. We propose that, before an accurate saccade, one of the saccade targets is selected, such that oculomotor cells centered on the saccaded location become more active in comparison to those encoding the non-saccaded target location. As our two targets were physically identical, saccade target selection probably occurs within the priority maps and propagates via top-down mechanism to the corresponding feature map columns (Baluch & Itti, 2011; Findlay & Walker, 1999; Marino, Trappenberg, Dorris, & Munoz, 2012; Trappenberg, Dorris, Munoz, & Klein, 2001). Then, within the brainstem, a winer-takes-all integration (Schiller et al., 1979; Scudder, Moschovakis, Karabelas, & Highstein, 1996a; 1996b) leads to an accurate saccade towards the selected target and the activity state within the feature map leads to higher sensitivity at the saccade endpoint before the eyes start to move (Figure 5B–C). Following the same rationale, we propose that averaging saccades follow from two equal activations within the priority and feature map columns associated with the saccade targets (Figure 5D).

**Figure 5.**
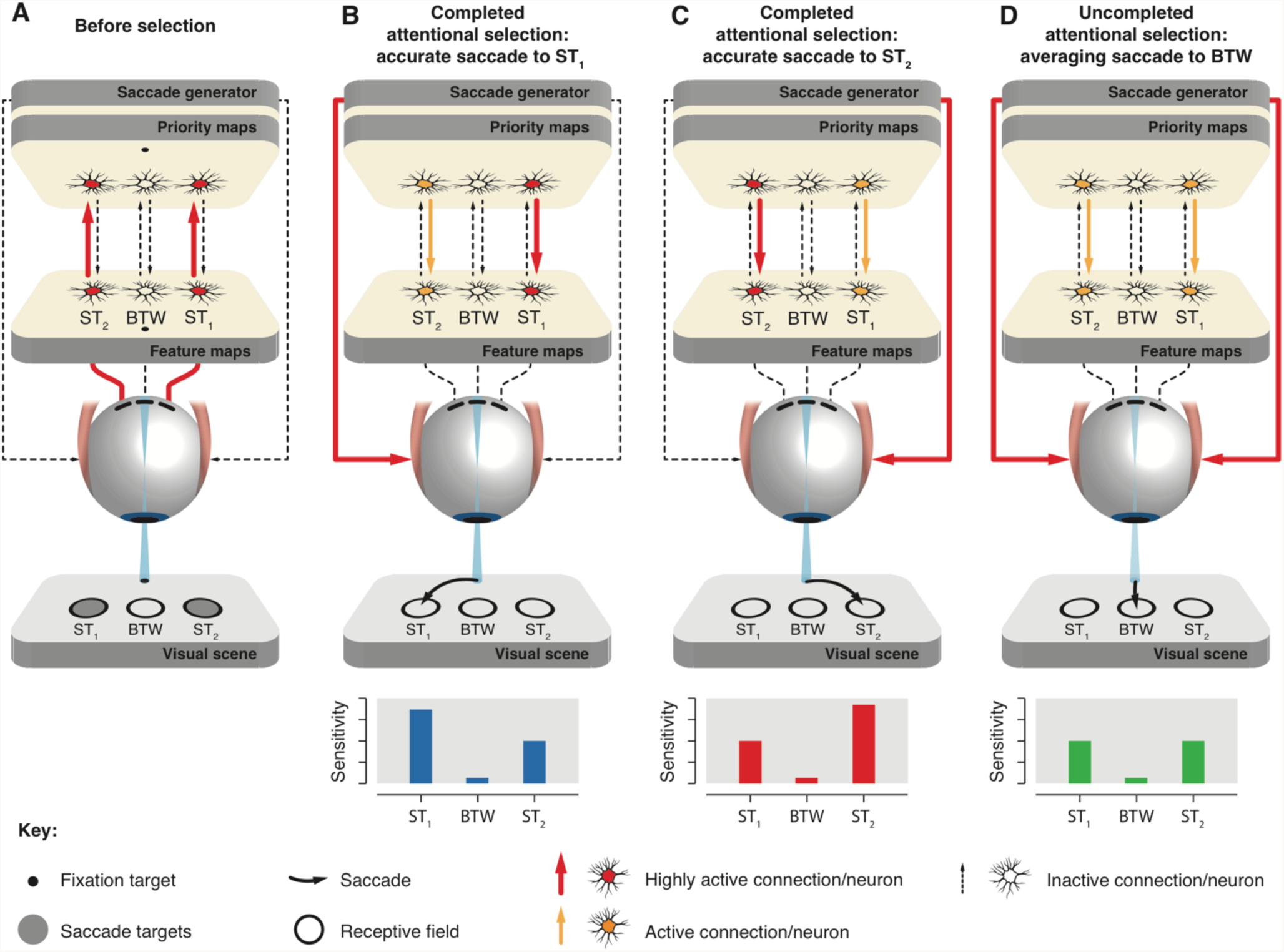
Attentional selection determines saccade endpoint. Two nearby saccade targets (gray dots) are flashed in the periphery from the fixation target and projected onto the retina, triggering a cascade of bottom-up (upward arrows) and top-down (downward arrows) processes throughout the visual processing hierarchy. Colors of the neurons and arrows indicate the level of activation. Each retinal projection connects to a specific neuron (in fact, a population of neurons) in retinotopic feature maps (V1–V4, MT). Feature map neurons, in turn, are linked to priority maps (FEF, LIP, SC). Priority maps activity is later integrated by a saccade generator (brainstem) driving the extra-oculomotor muscles. Note that the priority maps and the saccade generator are distinctive components within the processing hierarchy. Data panels show predicted sensitivity at the saccade targets (ST_1_ and ST_2_) and in between them (BTW) and black curved arrows show the predicted saccade path. **A.** Before attentional selection, ST_1_ and ST_2_ neuronal columns are highly activated by bottom-up connections driven by the saccade targets onset. **B–C.** Following a decay in the activity of both ST_1_ and ST_2_ neuronal columns, a completed attentional selection leads to a high activation of either ST_1_ (B) or ST_2_ (C) neurons in the priority maps, propagating via top-down connections to the feature maps. This leads to a pre-saccadic enhancement of sensitivity at the selected target and subsequently to an accurate saccade towards it. **D.** Uncompleted attentional selection leads to an equal and moderate pre-saccadic sensitivity enhancement at the saccade targets, but not in between, followed by an averaging saccade.

In other words, we propose that contrary to accurate saccades, selection is not completed at the onset of averaging saccades. Therefore, two equal signals representing the two saccade targets are integrated within the brainstem and trigger a saccade towards the intermediate location in space. Similarly, before saccade onset, the two saccade targets are associated with equal moderate attentional benefits. This proposal is supported by electrophysiological recordings showing that averaging saccades are associated with two distinct peaks within the intermediate layers of the SC (Edelman & Keller, 1998; Port & Wurtz, 2003). A similar, general conception of oculomotor programming was expressed by He and Kowler (1989), who proposed a two-stage process in which a single mechanism resolves attentional selection before the oculomotor program is being computed at a later stage based on attentional weighting. Our results and framework however go against a recent proposal that a merged activation within the superficial layers of the SC would feed back into the visual system (Vokoun et al., 2014) as this should have led to some attentional enhancement in between the saccade targets before an averaging saccade.

Our framework leads to some predictions with regard to the global effect. First, it predicts that any experimental manipulation modifying the difficulty of saccade target selection will directly impact the occurrence of averaging saccades. For example, specifying the location, the identity or the probability of a saccade target to appear at a certain location will decrease the task difficulty, thereby increasing the speed of the attentional selection process and reducing the occurrence of averaging saccades (Aagten-Murphy & Bays, 2017; Aitsebaomo & Bedell, 2000; Coëffé & O’Regan, 1987; He & Kowler, 1989; Heeman et al., 2014; Silvis et al., 2015). Also, it predicts that at a given latency, an easy saccade task should lead to fewer averaging saccades as compared to a more difficult one. Using a simple two saccade target task, it was shown that monkeys make averaging saccades only for express but not for normal saccade latencies (Chou et al., 1999), whereas they make averaging saccades even for normal saccade latencies in a task rendered harder by a visual search display (Arai et al., 2004). Similarly, Viswanathan and colleagues (2013) showed that at a saccade latency for which no consistent global effect was found with a distractor shown nearby a pro-saccade target, a clear global effect was evident with the same distractor shown nearby an anti-saccade target. These results are in line with our first prediction, as anti-saccades are associated with a slower attentional selection (Klapetek, Jonikaitis, & Deubel, 2016). Second, our framework predicts that one should not find any incremental presaccadic attentional benefit at one of the competing saccade targets before an averaging saccade, irrespective of the observed saccade latency. Future studies could directly test this prediction by measuring neuronal activity associated with the saccade targets before an averaging saccade. Third, we predict that motor cells with movement fields at the averaging saccade endpoint will stay silent before an averaging saccade, ruling out, at the neuronal level, the premotor theory of attention in a saccade task rather than in a covert attention task (Gregoriou et al., 2012; Ignashchenkova et al., 2004; Thompson et al., 2005).

Combining a measure of presaccadic visual sensitivity with a free choice saccade task we dissociated attention allocated to the intended saccade goal from attention allocated to the saccade endpoint. We report here that attention is not obligatorily coupled to an executed oculomotor program, ruling out the premotor theory of attention. Instead we propose that saccadic responses depend on the state of attentional selection at saccade onset.

## Materials and methods

### Participants

13 participants (age: 20-28, 7 females, 12 right-eye dominant, 1 author) completed the experiment for a compensation of 50€. The study was run over 2 experimental sessions (on different days) of 12 blocks of ~150 minutes each (including breaks). All participants except for one author (LW) were naive as to the purpose of the study and all had normal or corrected-to-normal vision. The experiments were undertaken with the understanding and written informed consent of all participants and were carried out in accordance with the Declaration of Helsinki. Experiments were designed according to the ethical requirement specified by the LMU München and with approval of the ethics board of the department.

### Setup

Participants sat in a quiet and dimly illuminated room, with their head positioned on a chin and fore-head rest. The experiment was controlled by an Apple iMac computer (Cupertino, CA, USA). Manual responses were recorded via a standard keyboard. The dominant eye’s gaze position was recorded and available online using an EyeLink 1000 Desktop Mount (SR Research, Osgoode, ON, Canada) at a sampling rate of 1 kHz. The experimental software controlling the display, the response collection as well as the eye tracking was implemented in Matlab (The MathWorks, Natick, MA, USA), using the Psychophysics (Brainard, 1997; Pelli, 1997) and EyeLink toolboxes (Cornelissen, Peters, & Palmer, 2002). Stimuli were presented at a viewing distance of 60 cm, on a 24-in Sony GDM F900 CRT screen (Tokyo, Japan) with a spatial resolution of 1,024 × 640 pixels and a vertical refresh rate of 120 Hz.

### Experimental design

Each trial began with participants fixating a central fixation target forming a black (~0 cd/m^2^) and white (~57 cd/m^2^) “bull’s eye” (0.4° radius) on a gray background (~19.5 cd/m^2^). When the participant’s gaze was detected within a 2.0° radius virtual circle centered on the fixation point for at least 200 ms, the trial began. At that time 24 distractor streams appeared equally distributed along a 10°-radius imaginary circle centered on the fixation target (see Figure 1A). Distractor streams consisted of flickering stimuli (40 Hz), alternating every 25 ms between a vertical Gabor patch (frequency: 2.5 cpd; 100% contrast; random phase selected each stream refresh; standard deviation of Gaussian window: 1.1°; mean luminance: ~28.5 cd/m^2^) and a Gaussian pixel noise mask (made of ~0.22°-width pixels with the same Gaussian envelope as the Gabors). After a random fixation period between 300 and 600 ms (in steps of 1 screen refresh: ~8ms), the fixation target switched off together with the onset of two saccade targets. Saccade targets, ST_1_ and ST_2_, were gray circles (~39 cd/m^2^; 1.1° radius, 0.2**°**-width) surrounding two randomly chosen streams with an inter-target angular distance of 30° or 90°. They were either presented transiently (50 ms) or continuously (until the end of the trial). Importantly, when presented transiently, the saccade targets had always disappeared from the screen at the time the discrimination target appeared on the screen. When presented continuously, on the other hand, the saccade targets always temporally overlapped with the presentation of the discrimination target. Our motivation to include these two saccade target durations was to check for a potential masking effect of the saccade targets on the discriminability of a discrimination target. Participants were instructed to select one of the saccade targets by moving their eyes towards it as fast and as accurately as possible. In 96.5% of all trials, between 75 and 175 ms after the saccade targets onset (a time determined to maximize discrimination target offset in the last 200 ms before the saccade), one of the 24 distractor streams was replaced by a discrimination target stream in which a tilted Gabor was played (25 ms, rotated clockwise or counter-clockwise by 12° relative to the vertical). The discrimination target could appear at any of the 24 distractor streams with equal probability and subjects were explicitly informed about this fact at the beginning of the experiment. In 3.5% of all the trials, we did not present any discrimination target, in order to evaluate its influence on saccade metrics (note that all other analyses are based on the discrimination target present trials). 500 ms after the saccade targets onset, all stimuli disappeared and participants were instructed to report the orientation of the discrimination target using the keyboard (right or left arrow key). Incorrect responses were followed by a negative-feedback sound. On trials in which no discrimination target was shown, participants’ response was followed by a random feedback sound.

3 participants were excluded from the analysis as their performance stayed at chance level irrespective of the position of the discrimination target. The remaining 10 participants completed between 6972 and 7055 trials of the saccade task. Correct fixation within a 2.0°-radius virtual circle centered on the fixation point was checked online. Trials with fixation breaks were repeated at the end of each block, together with trials during which a saccade started (i.e. crossed the virtual circle around the fixation target) within the first 50 ms or after more than 350 ms following the saccade targets onset (participants repeated between 46 to 395 trials across all blocks).

### Data pre-processing

Before proceeding to the analysis of the behavioral results we scanned offline the recorded eye-position data. Saccades were detected based on their velocity distribution (Engbert & Mergenthaler, 2006) using a moving average over twenty subsequent eye position samples. Saccade onset and offset was detected when the velocity exceeded or fell below the median of the moving average by 3 SDs for at least 20 ms. We included trials if a correct fixation was maintained within a 2.0° radius centered on the fixation target and if a correct saccade started at the fixation target and landed at a distance between 7° and 13° from the fixation target (±30 % of the instructed saccade size) and if no blink occurred during the trial. Finally, only trials in which the discrimination target offset was included in the last 200 ms preceding the saccade onset were included in the analysis (mean ± SEM discrimination target offset relative to the saccade onset for the selected trials: −50.2 ± 1.3 ms). In total we included 53117 trials in the analysis (78.2 % of the online selected trials; 75.7 % of all trials played) corresponding to an average of 106.0 ± 2.1 trials (115.9 ± 3.3 no discrimination target trials) and 105.3 ± 1.8 trials (125.0 ± 4.4 no discrimination target trials) per discrimination target location and participant, in the 90° and 30° condition, respectively.

Correctives saccades were defined as the saccades directly following the offline selected main saccades sequence and landing at a distance between 7° and 13° from the fixation target. Corrective saccades were included only if they started before the participant’s behavioral response and within the first 500 ms following the main saccade sequence. In total we obtained 14714 corrective saccade trials in the analysis (21.7 % of the online selected trials; 21.0 % of all trials played).

### Behavioral data analysis

Before proceeding to any behavioral analysis, we first rotated the trial configuration as to align the two saccade target locations (ST_1_: +15°, ST_2_: −15° and ST_1_: +45°, ST_2_: −45° for the conditions in which they were separated by 30° and 90°, respectively) symmetrically around the geometrical angle zero (BTW). We then determined the sensitivity to discriminate the orientation of the discrimination targets (d’): d’ = *z*(hit rate) − *z*(false alarm rate). To do so, we defined a clockwise response to a clockwise discrimination target (arbitrarily) as a hit and a clockwise response to a counter-clockwise discrimination target as a false alarm. Corrected performance of 99% and 1% were substituted if the observed proportion correct was equal to 100% or 0%, respectively. Performance below the chance level (50% or d’ = 0) were transformed to negative d’ values. We analyzed sensitivity as a function of the discrimination position in space irrespective of the saccade landing direction (Figure 3A-B) but also as a function of the discrimination target position relative to the saccade landing direction (Figure 3C–F). To do so, we redefined the position of the discrimination target relative to the saccade direction binned across 24 even angular sectors of 15° (±7.5° from each distractor stream center angle). This binning was chosen to match with the locations at which we tested visual attention.

We initially computed single subject means and then averaged these means across participants for each of the compared conditions to get the presented results. For all statistical comparisons we drew (with replacement) 10000 bootstrap samples from the original pair of compared values. We then calculated the difference of these bootstrapped samples and derived two-tailed *p* values from the distribution of these differences.

## Acknowledgments

The authors declare no competing financial interests.

